# MAVEN: Compound mechanism of action analysis and visualisation using transcriptomics and compound structure data in R/Shiny

**DOI:** 10.1101/2022.07.20.500792

**Authors:** Layla Hosseini-Gerami, Rosa Hernansaiz Ballesteros, Anika Liu, Howard Broughton, David Andrew Collier, Andreas Bender

## Abstract

**Background:** Understanding the mechanism of action (MoA) of a compound is an often challenging but equally crucial aspect of drug discovery that can help improve both its efficacy and safety. Computational methods to aid MoA elucidation usually either aim to predict direct drug targets, or attempt to understand modulated downstream pathways or signalling proteins. Such methods usually require extensive coding experience and results are often optimised for further computational processing, making them difficult for wet-lab scientists to perform, interpret and draw hypotheses from.

**Results:** To address this issue, we in this work present MAVEN (Mechanism of Action Visualisation and Enrichment), an R/Shiny app which allows for GUI-based prediction of drug targets based on chemical structure, combined with causal reasoning based on causal protein-protein interactions and transcriptomic perturbation signatures. The app computes a systems-level view of the mechanism of action of the input compound. This is visualised as a sub-network linking predicted or known targets to modulated transcription factors *via* inferred signalling proteins. The tool includes a selection of MsigDB gene set collections to perform pathway enrichment on the resulting network, and also allows for custom gene sets to be uploaded by the researcher. MAVEN is hence a user-friendly, flexible tool for researchers without extensive bioinformatics or cheminformatics knowledge to generate interpretable hypotheses of compound Mechanism of Action.

**Conclusions:** MAVEN is available as a fully open-source tool at https://github.com/laylagerami/MAVEN with options to install in a Docker or Singularity container. Full documentation, including a tutorial on example data, is available at https://laylagerami.github.io/MAVEN.

## 1 Background

The discovery of the mechanism of action (MoA) of a small molecule, which describes the biochemical interactions a molecule makes to produce a pharmacological effect, is an important aspect of drug discovery for a wide range of reasons, from repurposing for a new indication to anticipating potential side effects and rationalising phenotypic findings^1^. Advances in machine learning techniques, combined with large publicly availably bioactivity databases such as ChEMBL and PubChem, as well high-throughput biological assays such as LINCS L1000 and DRUG-Seq, have contributed to the development of computational methods for generating hypotheses of compound MoA^2^. Two popular approaches include target-based and network-based methods. Target-based methods aim to predict the direct biological target of the compound, and have shown high performance using chemical structure fingerprints as descriptors^3–5^. Network-based methods such as causal reasoning use transcriptomics data along with prior knowledge networks to infer upstream drivers of transcriptional changes, and have been shown to capture biological pathways modulated by drug compounds^6–9^.

However, such approaches often require proficiency in programming languages such as R and Python as well as the command-line, and output computer-readable data which can be difficult to convey to non-specialists, which can hinder scientific communication in multi-disciplinary groups. R/Shiny apps allow for the implementation of R code and the visualisation of results in an interactive GUI, and have been widely used, e.g., also for the integration of multi-omics (e.g., transcriptomics, phosphoproteomics, metabolomics) data with bioinformatics tools such as COSMOS^10^ and CARNIVAL^7^ to gain insights into compounds or other perturbations^11^. Hence, here we introduce MAVEN, or Mechanism of Action Visualisation and ENrichment, an R/Shiny app which allows users to integrate compound structure-based target prediction with gene expression-based causal reasoning without prior coding experience, and allows for the visualisation and pathway enrichment of the results to obtain a systems-level, easily interpretable view of the mechanism of action of a compound.

## 2 Implementation

MAVEN was written in the R programming language (v 4.2) using the Shiny application framework, and the source code is available for local installation at https://github.com/laylagerami/MAVEN. To run direct target prediction (which is optional for software functionality) the app also invokes PIDGINv4^4^ (https://github.com/BenderGroup/PIDGINv4) models and scripts implemented in Python, using a Bash command script called from within R. For causal reasoning over biological prior knowledge networks with CARNIVAL^7^ (https://github.com/saezlab/CARNIVAL), it is necessary to install an ILP (Integer Linear Programming) solver, either the free, open-source Cbc solver^12^ or the free-for-academic IBM ILOG CPLEX^13^. Installation and configuration instructions for the solvers are described in the documentation https://laylagerami.github.io/MAVEN/ along with troubleshooting steps. We also provide an R script to install all packages with the required versions, and a conda .yml file with packages required for running the PIDGINv4 Python scripts. In case a container solution is preferred, build files for Docker and Singularity containers with all required software and environments (including solvers) are provided. The Omnipath^14^ signed and directed protein-protein interaction network is included as well as gene expression^15^ and compound structure data for lapatinib which is used as an example in the documentation, and will be discussed here in the case study. For pathway enrichment on the predicted signalling network, MSigDB^16–18^ (v2022.1) gene sets in the hallmark (H), curated (C2) and ontology (C5) collections have been provided (as well as an option to use custom user-uploaded gene set files). Installation and deployment with the open-source Cbc solver have been tested on the HPC systems at Eli Lilly and Company and AWS in order to ensure compatibility with corporate computational environments.

The overall workflow for MAVEN is depicted in Figure 1. Three inputs are taken; known or hypothesised targets which can be predicted from the compound’s chemical structure with PIDGINv4^4^ or defined a priori (optional) (Figure 1A); a signed and directed (i.e., A activates/inhibits B) prior knowledge network (Figure 1B) for causal reasoning; and compound-induced gene expression data in the form of a summary statistic such as t-values or log2-fold changes (Figure 1C). A signed and directed prior knowledge network on causal protein-protein interactions is required to infer causality and function (activation or inhibition), and can be obtained from open source databases e.g., Omnipath^14^ (provided), SignaLink^22^ or SIGNOR^23^. Gene expression data in the form of differential expression signatures (i.e., Z-score, Log2FC, t-statistic) can be from any platform, e.g., microarray, RNA-Seq, and publicly available gene expression data is available for many perturbations in databases such as GEO (https://www.ncbi.nlm.nih.gov/geo/)^24^ (provided for the compound lapatinib) and LINCS L1000 (https://clue.io/releases/data-dashboard-Level5)^2^. The differential expression signature is then used to infer transcription factor (TF) activities with DoRothEA^20^ and pathway activities with PROGENy^21^, which is then used along with the prior knowledge network by CARNIVAL^7^ to optimise a subnetwork which captures signalling proteins upstream of TF activity changes and, if targets are predicted or provided, links them to the targets (Figure 1D). The outputs from DoRothEA, PROGENy and CARNIVAL are processed and formatted using helper scripts from (https://github.com/saezlab/transcriptutorial). Finally, the subnetwork can be viewed and exported to use in other software such as Cytoscape^25^, and we also provide a collection of MSigDB^26^ gene sets (or allow for the upload of a custom gene set) for pathway enrichment with over-representation analysis (ORA), the results of which can also be visualised on the network (Figure 1E).

**Figure 1:**
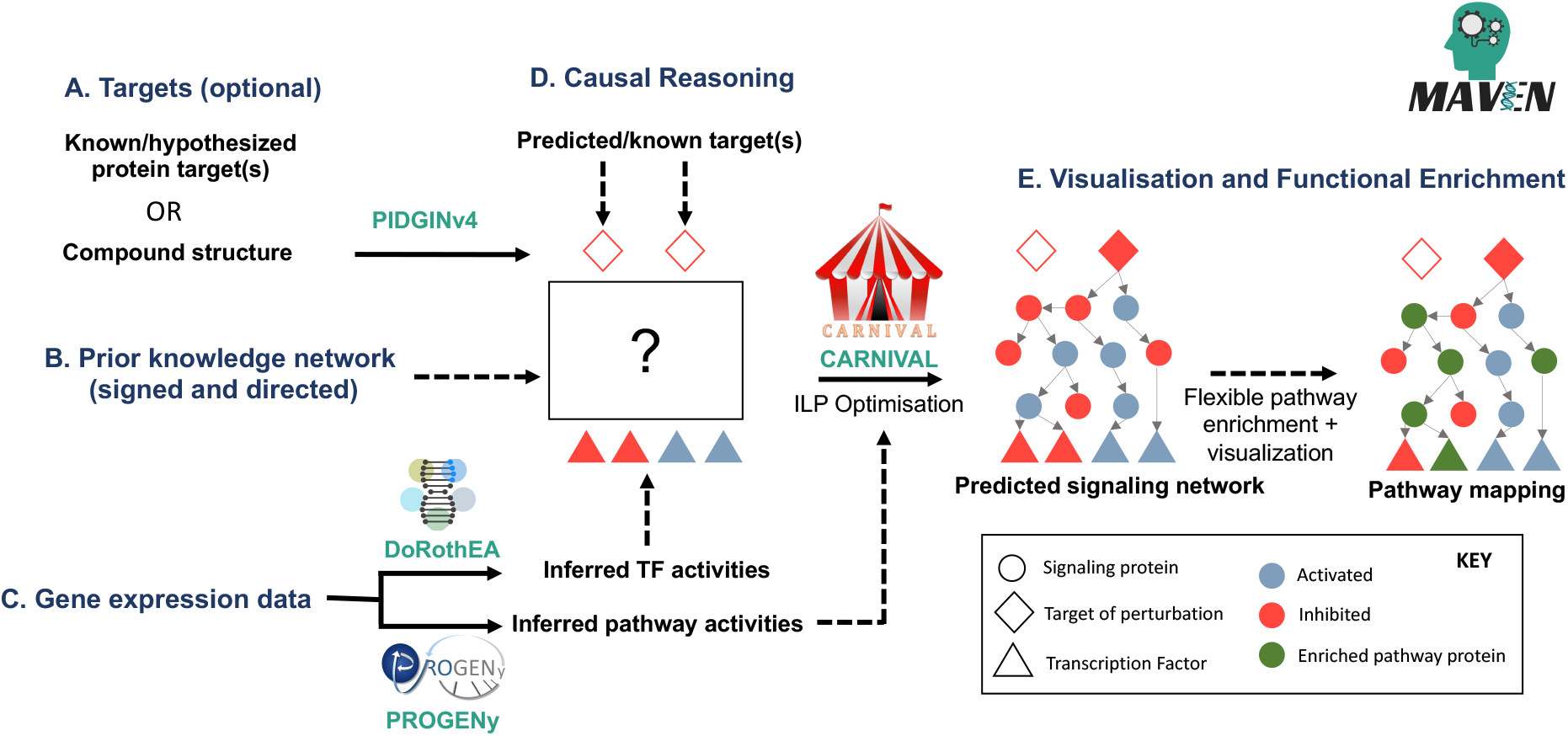
Workflow of analyses and features in MAVEN which requires 3 main inputs: Targets (A, optional) which can be known/hypothesised or predicted from the compound structure with PIDGINv4,^*4,19*^ a signed and directed prior knowledge network for causal reasoning (B) and differential gene expression signatures as e.g., log2FC, t-statistic (C) which is used for the inference of transcription factor and pathway activities with DoRothEA^*20*^ and PROGENy^*21*^. These are used as input to CARNIVAL^*7*^ for optimisation of the signalling network, connecting the targets to the TFs via intermediate signalling proteins (D). The resulting predicted signalling network is displayed in the GUI and interactive pathway enrichment can be carried out to contextualise the signalling proteins and interpret the compound’s mechanism of action (E). Figure adapted with permission from https://github.com/saezlab/carnival.

Throughout the workflow, all chosen parameters and command-line options are saved in log files for reproducibility purposes and so that analyses can be re-run programmatically (for information on how to run the tools used in MAVEN, please refer to https://pidginv4.readthedocs.io/en/latest, https://github.com/saezlab/shinyfunki and https://github.com/saezlab/transcriptutorial). Additionally, there are help buttons throughout the GUI with more information to aid the user in choosing algorithm parameters, and guidelines on the formatting of data.

The functionalities implemented in MAVEN will now be discussed in more detail:

### Target prediction with PIDGINv4

The first data analysis step in MAVEN is target prediction based on compound chemical structure - though this is optional and targets can be manually entered or left out entirely. These targets are used as input to CARNIVAL in a later step, to connect to inferred signalling proteins. Target prediction is implemented in MAVEN by invoking the PIDGINv4 software. PIDGINv4^4^ is an open-source target prediction tool trained on ChEMBL^27^ (v29) and PubChem^28^ data using the scikit-learn^29^ Python package, available on GitHub (v4.2). The tool consists of a collection of Random Forest models trained on the chemical structures (ECFP4 fingerprints calculated with the RDKit^30^ Python package) of active and inactive compounds against 2,000+ human targets, and Python scripts to generate predictions for query compounds and to search for structurally similar compounds in the model training sets. For target prediction, the user is required to upload a .smi file, and a ChemDoodle^31^ widget^32^ is embedded in the app GUI to sketch the structure and generate a SMILES file in case the structural SMILES are not known. The user can select various parameters for the target prediction including activity threshold (0.1, 1, 10 or 100 *μ*M – default 10 *μ*M), number of cores (default 10), and applicability domain filter (default 50 out of 100) to remove low-confidence predictions^33^. Once the user chooses to run the target prediction, a Bash script is invoked which runs the predict.py and sim_to_train.py PIDGINv4 scripts. The predict.py script processes the input SMILES and calculates ECFP4 fingerprints, applies the pre-trained models, and then outputs the Platt-scaled Random Forest probability values. The sim_to_train.py script retrieves the most structurally similar compound in the ChEMBL29 database (nearest neighbour), based on Tanimoto similarity of their ECFP4 fingerprints. The results from both scripts are saved on disk, and then formatted and displayed in the GUI.

### Transcription factor enrichment with DoRothEA and VIPER

To perform causal reasoning on a protein-protein interaction network, the gene expression data must be converted from the “gene-level” to the “protein-level” by inferring upstream TFs driving the expression changes. DoRothEA (Bioconductor dorothea v1.8.0) describes curated TF regulons, so known TF-gene interactions^20^. Each interaction is given a confidence score reflecting the supporting evidence behind it from A (highest confidence, manually curated) to E (lowest confidence, computational predictions). The package is coupled with the VIPER (v1.3)^34^ statistical method to infer TF activity from gene expression data, generating normalised enrichment scores for each TF^35^. In the app, the user can select the confidence levels A-E to filter the interactions in the regulon, and a slider for the number of top TFs they want to report and use for causal reasoning analysis. By default, 50 TFs are reported and plotted as a bar-chart in terms of their normalised enrichment score (NES, from −1 to 1). This number is generally a trade-off between coverage and noise which can be examined by adjusting the slider and viewing the NES plot, which updates automatically upon re-calculation. Furthermore, only confidence levels A-C are included by default, but this criterion can be relaxed if more enriched TFs are required. The documentation and help buttons also provide guidance on choosing these parameters. Another parameter which can be changed in the source code (but not the GUI) is the ‘*minsize*’ VIPER parameter which indicates the minimum number of genes per TF regulon, set to 5 by default.

### Pathway activity inference with PROGENy

Pre-weighting proteins on the prior knowledge network has shown to improve the causal reasoning results by CARNIVAL^7^. PROGENy^21^ (Bioconductor progeny v1.16.0) is a “footprint” method which infers pathway activities by leveraging a large compendium of publicly available perturbation experiments that yield a common core of Pathway RespOnsive GENes. Based on the pathway footprint genes, PROGENy produces pathway scores for 14 major signalling pathways (Androgen, EGFR, Estrogen, Hypoxia, JAK-STAT, MAPK, NFkB, p53, PI3K, TGFb, TNFa, Trail, VEGF and WNT). The pathway scores are converted under-the-hood to protein weights to improve the CARNIVAL optimisation. The user can select the number of most dysregulated genes to include in the PROGENy pathway score calculations (by default 100, but this depends on the number of input genes – for experiments with a higher coverage such as RNA-Seq, this can be increased to e.g., 200 – 500).

### Causal reasoning with CARNIVAL

The last analysis tool implemented in MAVEN is causal reasoning to infer dysregulated signalling networks. CARNIVAL^7^ (Bioconductor CARNIVAL v.2.6.2) is a causal reasoning algorithm based on integer linear programming (ILP) which aims to optimise a subnetwork of signalling proteins contextualising a perturbation of interest. CARNIVAL takes as input dysregulated transcription factors (from DoRothEA) and a prior knowledge network (signed and directed protein-protein interactions), with pre-computed node weights (based on pathway activity scores from PROGENy) to aid the network optimisation. CARNIVAL generates multiple solutions which are then aggregated to form a consensus network which connects TFs to targets (pre-defined or predicted with PIDGINv4) *via* inferred dysregulated signalling proteins, including their sign (activated or inhibited). If no target is defined then the signalling proteins get connected to a proxy “perturbation” node. The user can choose the runtime (in seconds) and the number of cores. To solve the ILP problem a separate solver must be installed - Cbc^12^ (v.2.9, free and open source) or IBM ILOG CPLEX^13^ optimisation studio (v20.10, free for academic use or a license is required). The lpSolve (v5.7.16) ILP optimiser^36^ implemented in R is also available to use and is installed along with CARNIVAL, but it is strongly recommended to be used only for toy examples or testing purposes.

### Visualisation and functional enrichment

Following causal reasoning with CARNIVAL the consensus network is visualised in the GUI using the visNetwork package (CRAN visNetwork v.2.1.0). To put the inferred signalling network into biological context it is possible to perform functional enrichment. To this end, 11 MSigDB^26^ gene sets collections are included with MAVEN (such as Hallmark^16^, GO^18^, Reactome^37^, Wikipathways^38^). Alternatively, a .gmt file can be uploaded by the user for custom enrichment analysis. Over-representation analysis of the signalling network nodes in the gene sets using the prior knowledge network as background is performed with piano^39^ runGSAhyper function (Bioconductor piano v2.10.1). Following enrichment and tabular display of the results, the user can select a pathway of interest and highlight the participating proteins on the network. The pathway results can also be downloaded. The network .sif file is also saved for further analysis and visualisation in Cytoscape^25^ or other software packages.

## 3 Demonstration and Discussion

To demonstrate the app’s utility for generating hypotheses for compound mechanism of action in practice, and to give an overview of the UI and app functionalities, we will now present a case study using the EGFR and ERBB2 (HER2) inhibitor lapatinib (we also provide this as a tutorial included in the documentation).

The differential gene expression data used in this case study is derived from lapatinib-treated (1uM, 6h) HER2-positive BT474 breast cancer cells, from a publication by Sun et al^**15**^ (GEO^**24**^ accession GSE129254). In HER2-positive breast cancer, lapatinib inhibits the activation of signalling pathways downstream of EGFR and HER2 including MAPK, PI3K-AKT and PLC-γ, leading to apoptosis, decreased cellular proliferation and cell cycle arrest^**40**^. The aim of the MAVEN analysis in this case study is to infer a signalling network which captures the known cellular response of HER2+ positive cells treated with lapatinib.

The MAVEN UI is split into five tabs; Index (landing page), Data, Targets, Analysis and Visualisation (Figure 2). The landing page provides a summary of the MAVEN workflow and the case study will proceed from the second tab (Data).

**Figure 2:**
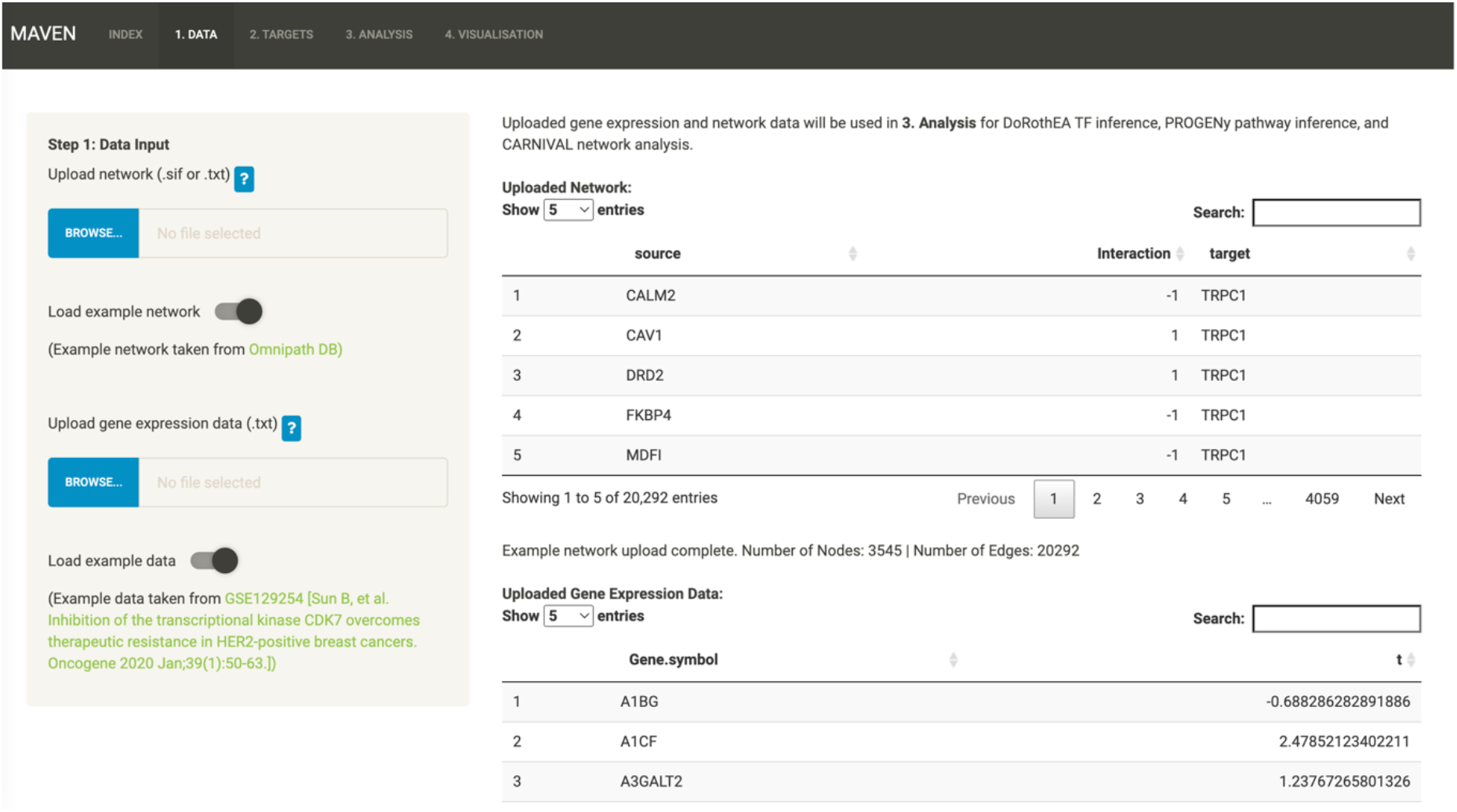
Screenshot of the Data tab showing the overall layout of MAVEN’s GUI after the case study datasets were loaded into the app via the “Load example” toggles

### Data

Here the gene expression data and prior knowledge are uploaded and stored in local memory for use in the Analysis tab. The user can browse for their files or use the toggle to load the Omnipath network and the lapatinib gene expression data used in this case study (Figure 2). As well as the documentation, there are help buttons throughout the workflow to explain file formats, definitions of parameters, and so on.

After checking that the data is in the correct format (including checking valid HGNC symbols and reporting any invalid symbols using HGNChelper v0.8.1^**41**^), the GUI provides a summary of the uploaded data for the user to check e.g., number of nodes and edges in the network. The user is then prompted to move onto the Targets tab.

### Targets

The Targets page is split into four sub-tabs (Figure 3) and is an optional step in the MAVEN workflow. In the first tab (Figure 3A), the user either uploads a SMILES file or sketches their compound to produce a SMILES file. Following successful SMILES upload, the compound is displayed as an image for the user to check, which can be seen for the case study with the correctly rendered lapatinib structure. In the second tab, the user is able to select the options for running PIDGIN (Figure 3B). Here, the bioactivity threshold was set to 1*μ*M to correspond with the concentration of lapatinib used to generate the gene expression data. The applicability domain (AD) filter was set to 30, and 20 cores of compute power were used to run the predictions. After choosing the parameters the user is prompted to browse for the location of their PIDGINv4 installation directory, and then a button becomes available to click for running the target prediction analysis (which can be monitored *via* the R console output). Targets can also be defined manually by entering their HGNC symbols in the (D) User-defined targets tab.

**Figure 3:**
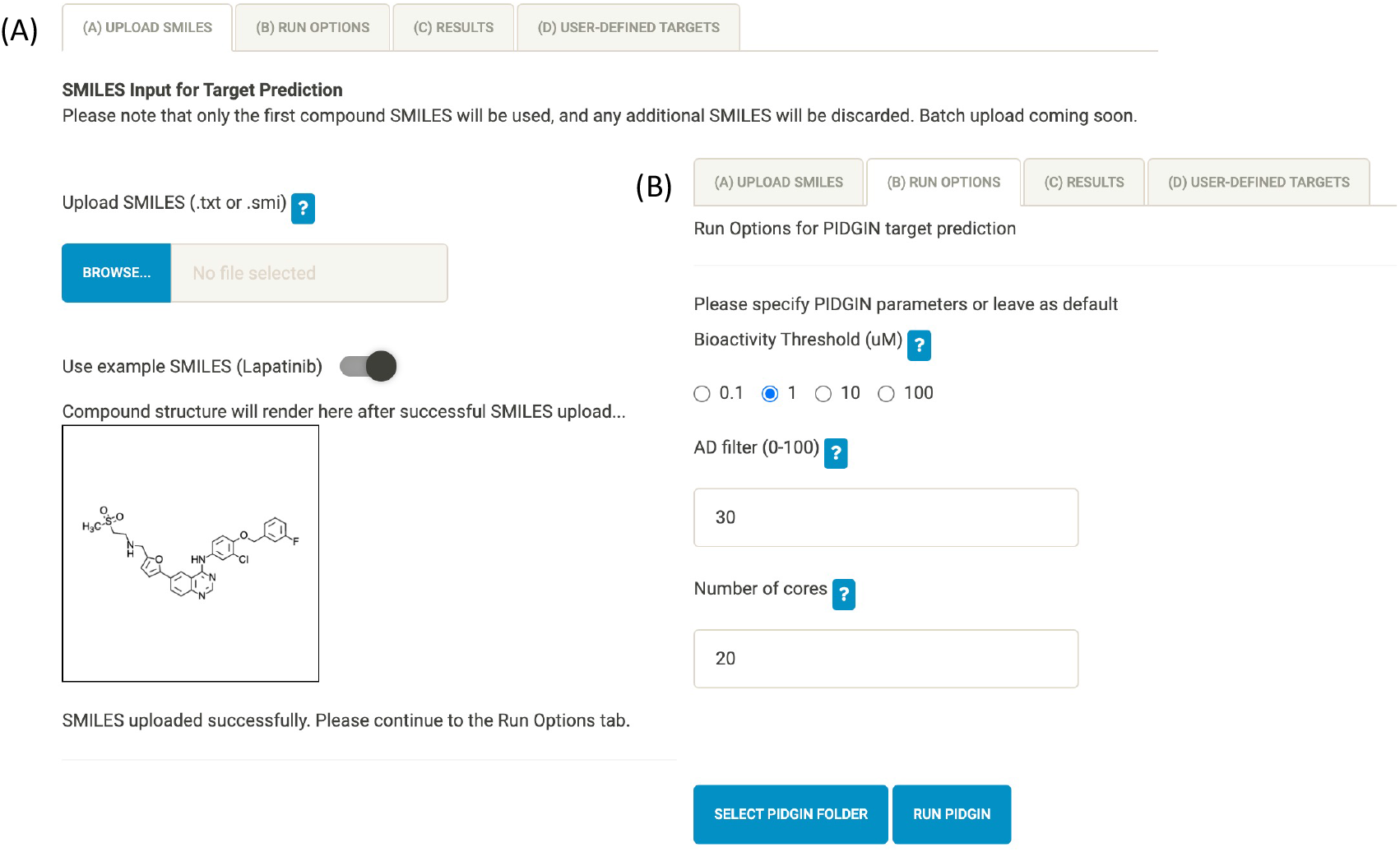
The Targets page contains 4 sub-tabs including SMILES upload (A) and PIDGIN run options (B). Results are displayed in (C) Results, and the user can manually define targets in (D) User-defined targets (not shown)

Once the PIDGIN run is complete, the results are saved and also displayed in the third tab, (C) Results, as a data table (Figure 4), with one row for each target model. The table contains the HGNC symbol, target name, predicted probability of activity, ChEMBL ID of the most structurally similar compound in the ChEMBL29 database (nearest neighbour), Tanimoto similarity of the nearest neighbour compared to the query compound computed from ECFP4 fingerprints, and the experimental measurements available for this compound. The target and nearest neighbour are hyperlinked to the UniProt and ChEMBL databases, respectively. It can be seen that many of the highest-predicted targets (ERBB4, EGFR, ERBB2, KCNH2 and PIK3C2B) are experimentally measured targets of lapatinib (CHEMBL554, Tanimoto Similarity = 1).

**Figure 4:**
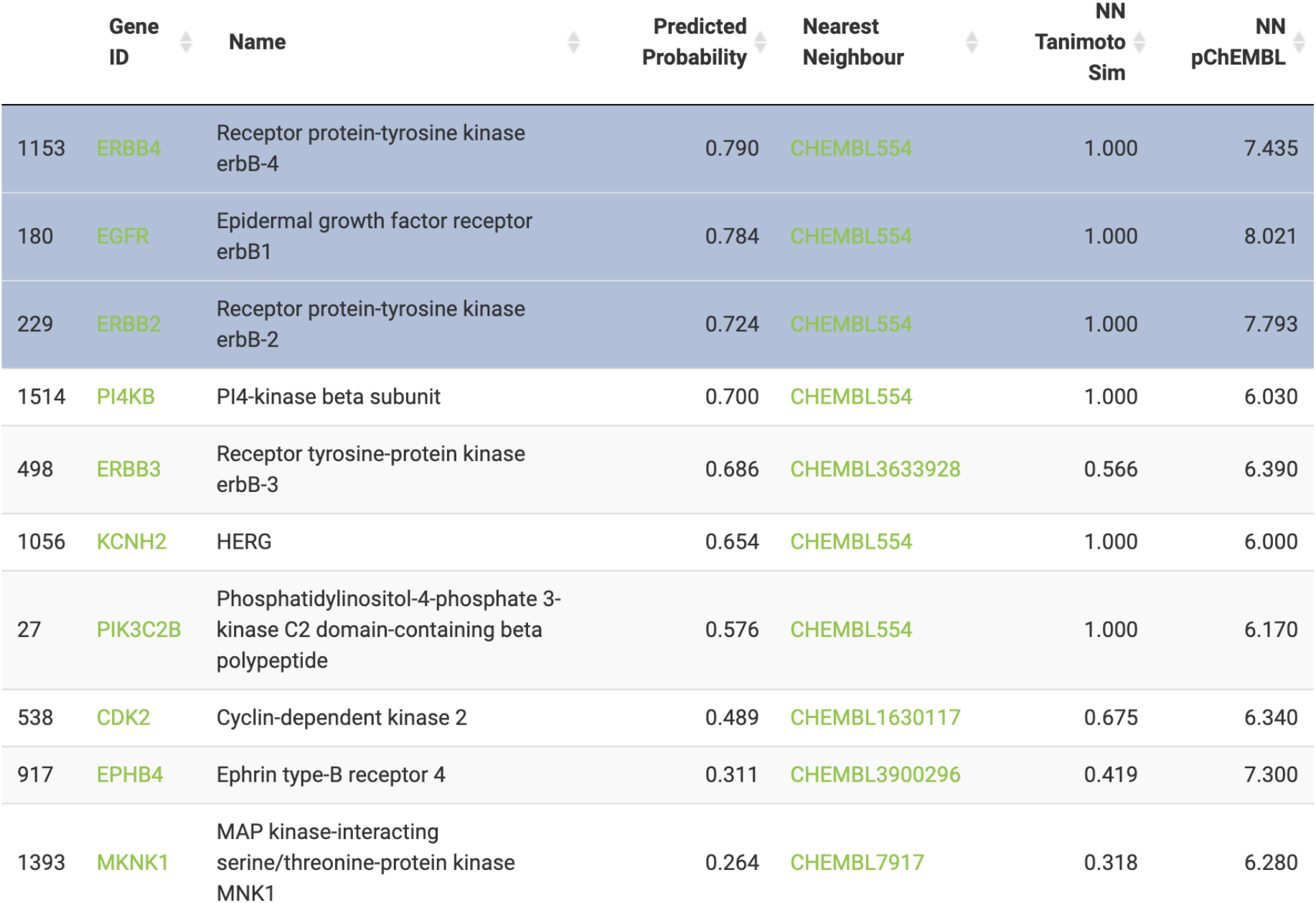
Loaded data table of PIDGIN results. The results table includes a row for each target defined by their HGNC symbol (hyperlinked to the UniProt database entry) and preferred names, the predicted probability of activity of the query compound lapatinib, the most structurally similar compound (nearest neighbour - NN) in the model training set (hyperlinked to the ChEMBL database entry), the Tanimoto similarity of the NN compared to lapatinib (where 1 indicates the compounds are exactly the same), and the experimental pChEMBL value of the NN against the target. Here the top three predicted targets (ERBB4, EGFR and ERBB2/HER2), are selected for further analysis to recapitulate lapatinib’s MoA in HER2+ cells.

Targets can be chosen from the PIDGIN output (by selecting rows) based on the predicted probabilities as well as Tanimoto similarities (the higher the better in both cases), or by consulting the literature references to a wide variety of protein functions listed in their linked UniProt entries (e.g., https://www.uniprot.org/uniprot/P00533 for EGFR). Alternatively, the analysis can be run without targets, and then re-run with selected targets based on these findings to investigate specific target hypotheses. For example, if the final network outputs nodes from a particular signalling pathway, a highly-predicted target upstream of this pathway can be used to refine the final network. The information provided in Figure 4 is intended only for selecting targets of interest, only the target HGNC symbols themselves are used as information for the CARNIVAL optimisation.

We took the three targets with highest predicted probability; ERBB4 (0.790), EGFR (0.784) and ERBB2/HER2 (0.724) all known to be expressed in HER2+ breast cancer^**42**^^**43**^, to the causal reasoning analysis stage. The rows are selected in the data table as shown in Figure 4.

### Analysis

The analysis page is split into three sub-tabs for the three bioinformatics analysis methodologies; DoRothEA, PROGENy and CARNIVAL. The settings for each can be set on the left-hand side of the page (Figure 5).

**Figure 5:**
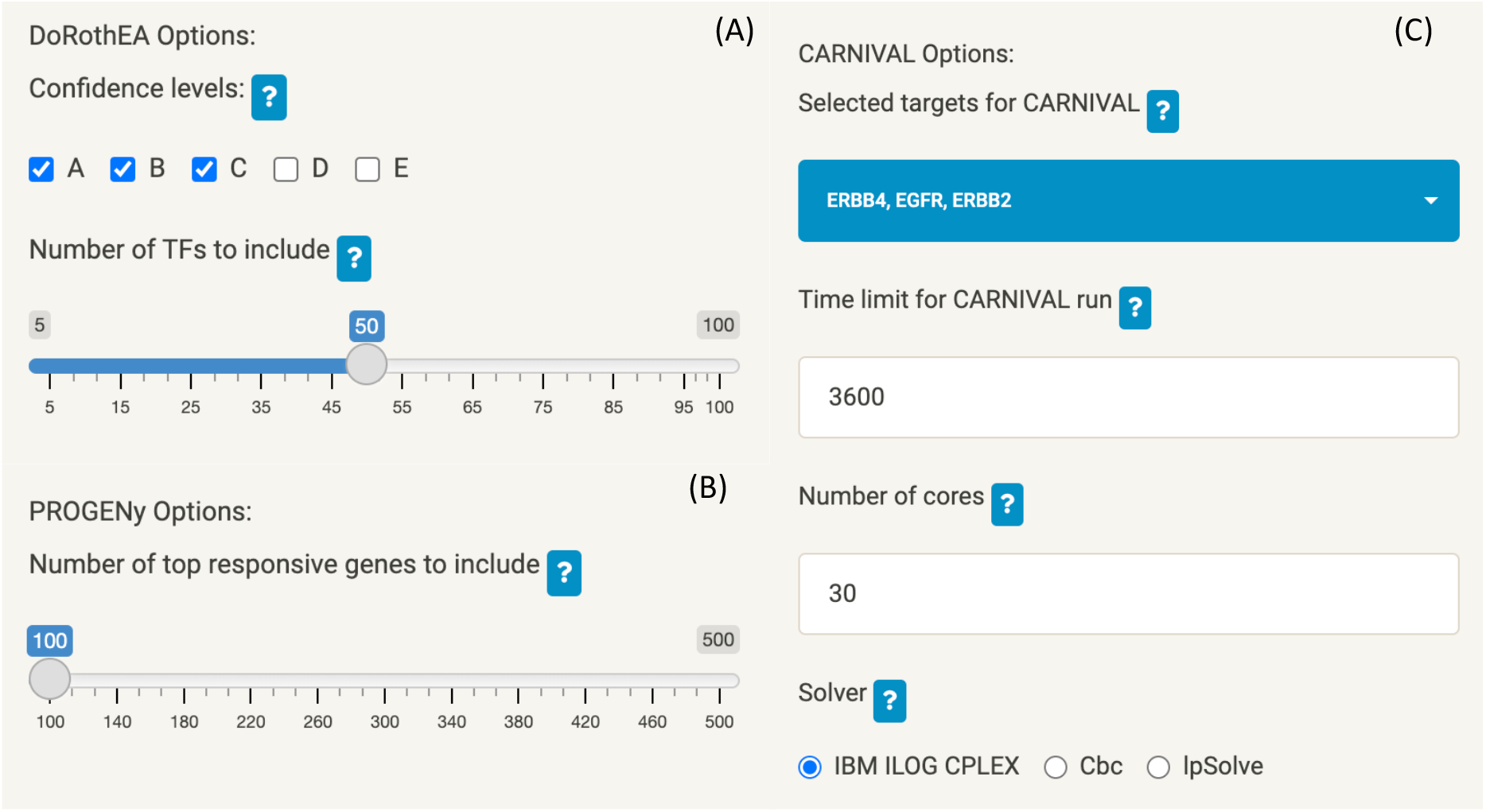
Settings for DoRothEA (A), PROGENy (B) and CARNIVAL (C) which can be interactively set in the MAVEN GUI

For the case study, DoRothEA (Figure 5A) was run with confidence levels A, B and C and the top 50 enriched TFs have been used for further analysis. For PROGENy (Figure 5B) the top 100 most responsive lapatinib genes (based on the t-values input in the Data stage) were used for the calculations. For CARNIVAL (Figure 5C) it can be seen that the targets selected in the previous step (Figure 4) have populated the CARNIVAL options, and they can be further deselected if required - here, we kept EGFR, ERBB4 and ERBB2 as described above. We set a time limit of 3,600 seconds for the calculations, 30 cores of compute power and used the IBM ILOG CPLEX solver for solving the ILP problem. This means that the solver will generate as many optimal network solutions as possible with the given time and compute resources, and output the final consensus network. Increasing the time limit or number of cores hence allows the solver to generate more networks, which may be required if no optimal solutions are found.

Following DoRothEA analysis, the resulting normalised enrichment scores (NES) for each TF are displayed as a bar chart (Figure 6) and a corresponding data table with TFs hyperlinked to their corresponding UniProt page. It can be seen from the plot that the top enriched upregulated TF was FOXO3 which is known to be upregulated by lapatinib in HER2+ cells^**44**^, and the top enriched downregulated TF was ESRRA which is known to be degraded in response to lapatinib-mediated inhibition of growth factor-induced signalling in HER2+ tumours^**45**^. Hence, MAVEN is able to generate an easy-to-interpret overview of TFs which are known to be dysregulated by lapatinib in the specific cellular context under investigation.

**Figure 6:**
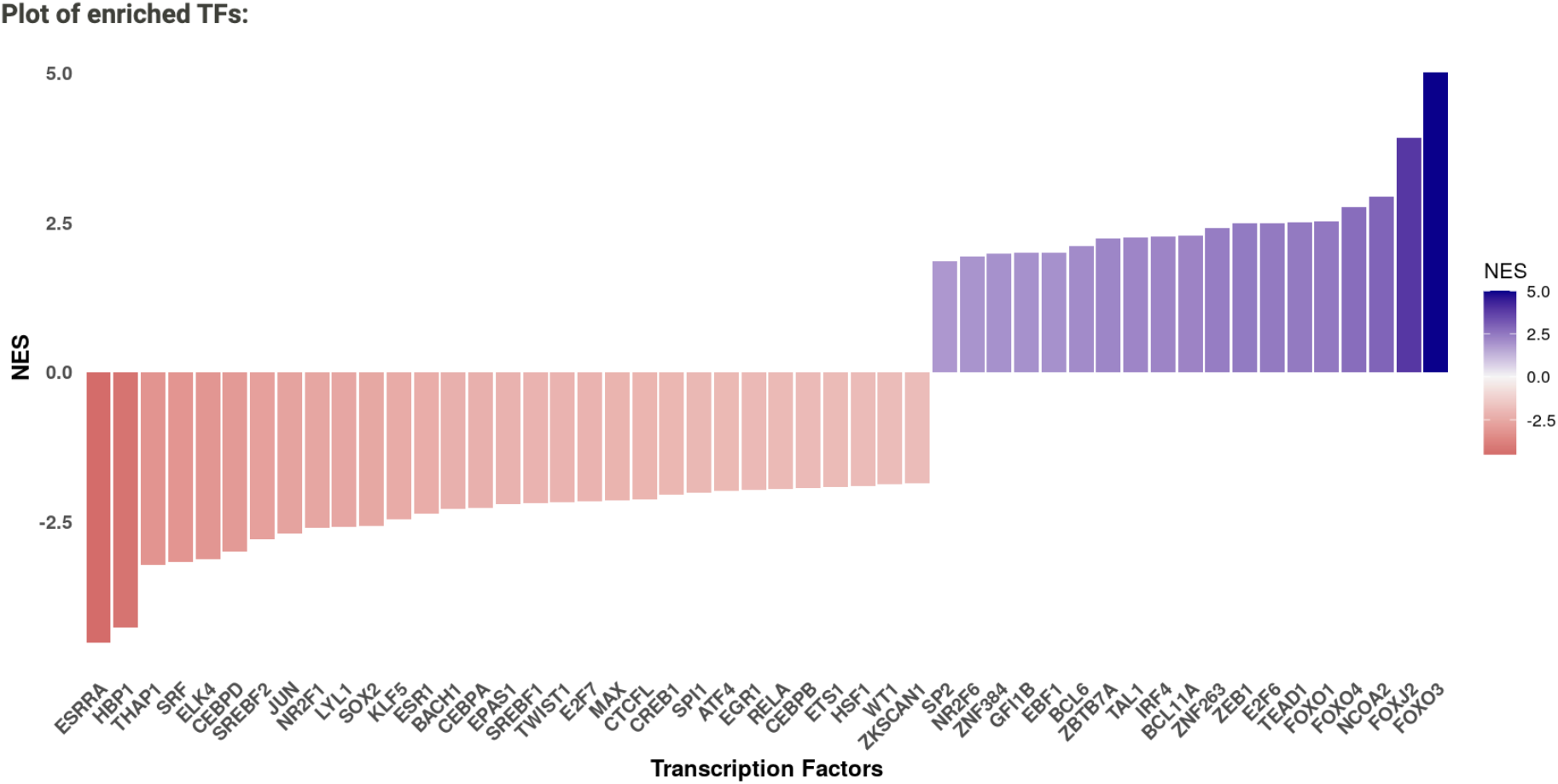
DoRothEA results derived from the differential expression signature of lapatinib-treated HER2+ BT474 cells expressed as a colour-coded bar plot. NES = Normalised enrichment score. Here, the most enriched upregulated TF (indicated by a positive NES) was FOXO3, and the most enriched downregulated TF (indicated by a negative NES) was ESRRA, which matches with the known activity of lapatinib in HER2+ cells.

If the slider is adjusted to select a different number of top-scoring TFs, the plot and table of results automatically update. The number chosen here is a trade-off between coverage (where selecting a higher number may lead to additional findings) and also noise, where on the other hand a greater number of TFs may not necessarily contribute additional information and instead increase computational time. To aid in this decision, the plot and associated UniProt information for each TF can be consulted to select a number that provides good coverage of different protein functions (i.e., to not solely choose a set of proteins in the same family, so that the CARNIVAL analysis can better exploit the prior knowledge network) coupled with prior knowledge/hypotheses on phenotypic findings. The interface help buttons (which can be seen in Figure 5) also provide guidance text for selecting these parameters, from the authors of DoRothEA.

Following PROGENy analysis, the results are visualised in the same way - a bar chart of predicted pathway activity score (from −1 to 1 indicating inhibition and activation) (Figure 7) and a corresponding data table (not shown). In agreement with the results of the analysis, lapatinib is known to inhibit the EGFR^**46**^, MAPK^**47**^ and PI3K^**48**^ pathways in HER2+ cells. The pathway scores are converted to weights on the protein-protein interaction network, which aids the optimisation of the signalling subnetwork by CARNIVAL^**7**^. By default, the top 100 top responsive genes are chosen, but this can be adjusted depending on the coverage of the gene expression experiment – in general, the greater the number of genes measured, the greater the number of top responsive genes (e.g., 200-500 for RNA-Seq experiments). The bar chart will again update upon adjustment of the number of genes, and can be interpreted with regards to the function of each pathway and what would be expected based on what is known about the compound.

**Figure 7:**
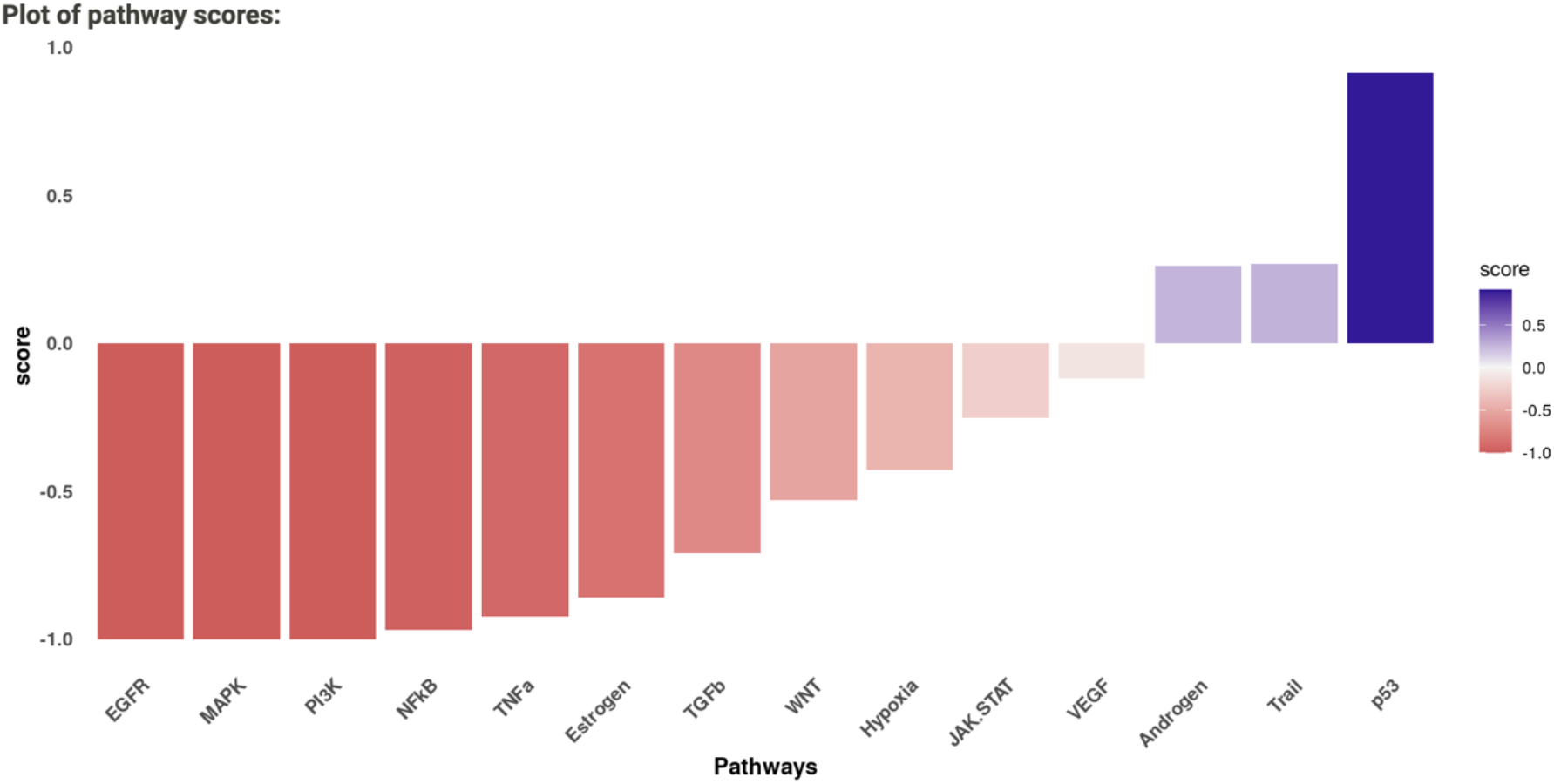
PROGEny results for lapatinib-treated differential expression signature in terms of predicted pathway activity score (from −1 to 1 indicating inhibition and activation). It can be seen that known pathways inhibited by lapatinib in HER2+ cells EGFR, MAPK and PI3K, are predicted as inhibited, aiding the optimisation of the CARNIVAL network in the next stage of the analysis.

### Visualisation and enrichment analysis

Following DoRothEA and PROGENy analysis, CARNIVAL can be run, taking as input the DoRothEA enriched TFs and pathway weights from PROGENy, as well as the prior knowledge network uploaded in the first Data step. Once complete, the resulting CARNIVAL consensus network is visualised on the visualisation page (Figure 8). Files from previous analysis runs, which are automatically saved, can also be uploaded into the tool for visualisation.

**Figure 8:**
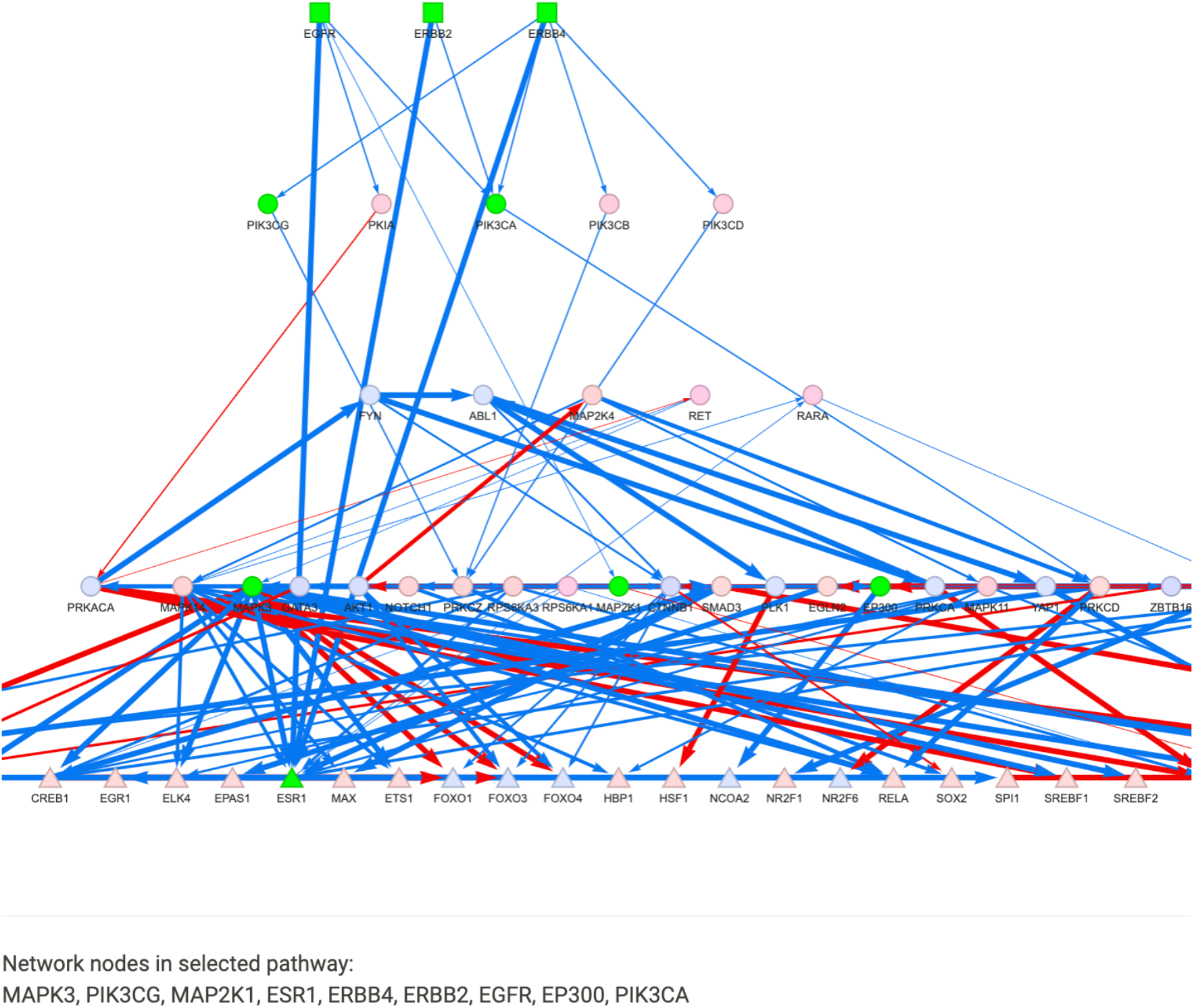
Following CARNIVAL analysis (or re-uploading previous results) the user can carry out pathway enrichment and visualisation on the inferred signalling network. Here, pathway enrichment was carried out on the network derived from lapatinib data with the built-in Biocarta MSigDB gene set, and the HER2 signalling pathway is highlighted on the network (as green nodes).

It can be seen that the top layer of the network consists of the three selected targets (ERBB4, ERBB2 and EGFR), the bottom layer consists of the input TFs (e.g., FOXO1, FOXO3, ESR1), and they are connected by signalling proteins with inferred directionality (indicated with blue for up-regulation and red for down-regulation), which along with their interactions have been optimised from the input prior knowledge network. As well as visualising the resulting network and deriving hypotheses from individual nodes, it is possible to perform pathway enrichment using the network proteins in an over-representation analysis. To illustrate this, we ran the enrichment analysis using the BioCarta^**49**^ gene set and the top enriched pathway, HER2 signalling pathway (adjusted p-value = 2.26e-9) is visualised on the network with participating proteins highlighted in green (Figure 8). Hence, CARNIVAL was able to construct a signalling network highly enriched for HER2 signalling, including the signalling proteins MAPKs, ESR1, ERBB2, EGFR, PIK3CA^**50**^ and EP300^**51**^, which are known to be relevant for the primary mechanism of action of lapatinib in HER2+ cancers^**50**^.

The enrichment results are also displayed in the GUI as a data table (Figure 9) and if one of the included MSigDB sets was used for the analysis, then they can be clicked through to the entry on the MSigDB website. A .csv file with more information on the enrichment results (e.g., participating proteins, odd’s ratio, unadjusted p-value) can also be downloaded.

**Figure 9:**
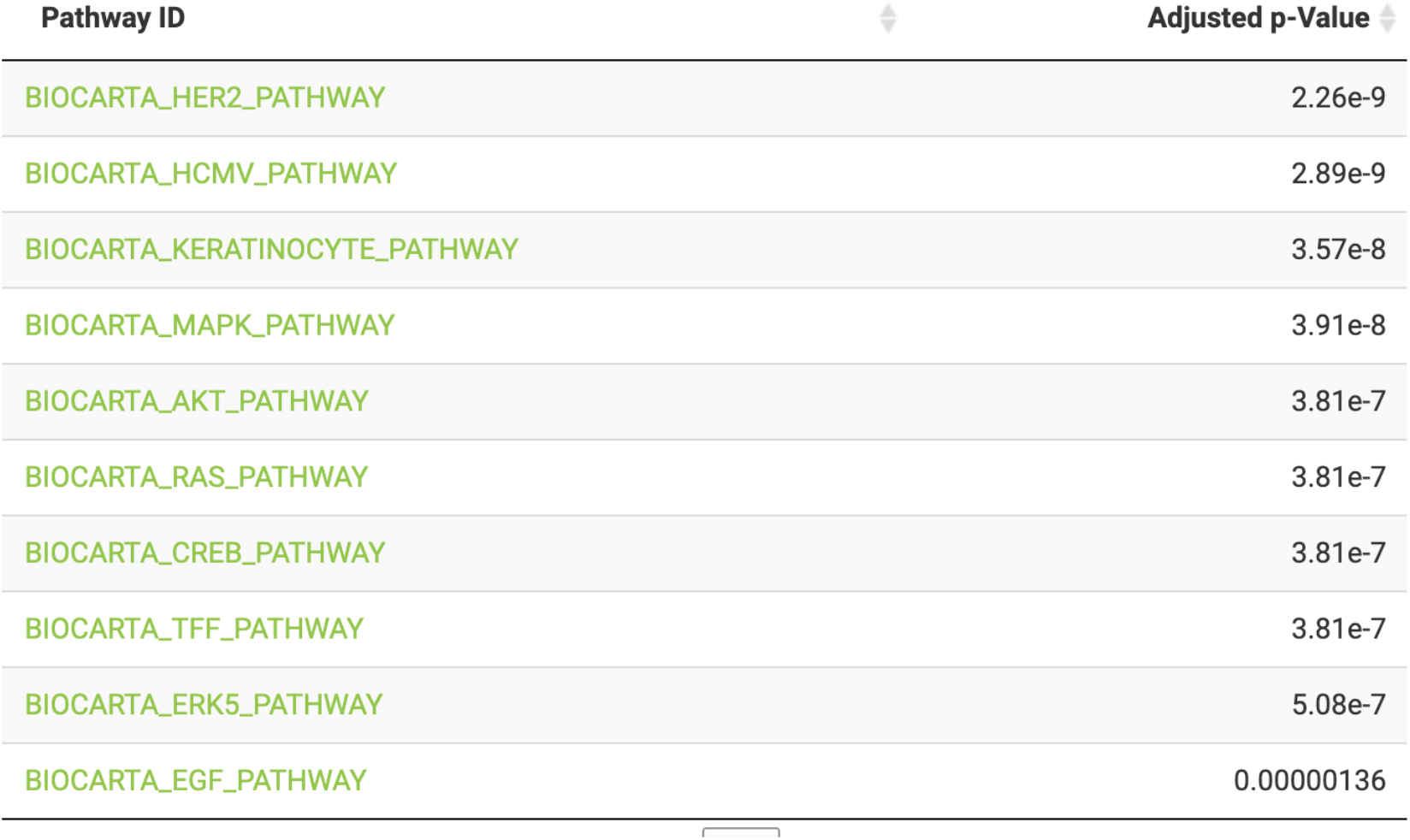
Pathway enrichment results table following over-representation analysis on the lapatinib-derived signalling network. It can be seen that the HER2 pathway is the top enriched pathway; selecting it lights up participating on the signalling network and reports the list of proteins to the user. The full results can also be downloaded as a .csv file.

### Case Study Summary

Through the case study, we have demonstrated the ability of the MAVEN R/Shiny app and its constituent tools to produce and report correct target prediction results (predicting the lapatinib targets EGFR and ERBB2), infer both down- and up-regulated transcription factors induced by lapatinib (including FOXO3 and ESRRA), infer pathways known to be modulated by lapatinib (EGFR, MAPK and PI3K), and finally construct and visualise a signalling network which is highly enriched for the HER2 signalling pathway known to be modulated by lapatinib. This demonstrates the detailed insights into compound MoA that can be obtained using MAVEN’s user-friendly interface, and without requiring extensive coding knowledge.

### Future Development

Future additions to the app will include a batch upload option to analyse multiple compounds at once and compare their results, options to use other causal reasoning algorithms, and the ability to upload and analyse other data types (e.g., phosphoproteomics and metabolomics data). Supplementary files such as the MSigDB gene sets will be continuously updated to reflect any major version changes. Suggestions for new features can also be requested on the GitHub page https://github.com/laylagerami/MAVEN.

## 4. Conclusions

We have developed an R/Shiny app called MAVEN (Mechanism of Action Visualisation and ENrichment) a novel, feature-rich tool integrating chemical-structure-based target prediction with gene expression-based causal reasoning analysis, coupled with visualisation and pathway enrichment analysis. A case study, using the chemical structure of lapatinib and gene expression data derived from lapatinib-treated HER2+ positive cells, has demonstrated the ease of inferring detailed insights into compound MoA using MAVEN.

## Availability and requirements

**Project name:** MAVEN

**Project home page:** https://github.com/laylagerami/MAVEN

**Operating system(s):** Platform independent

**Programming language:** R, Python

**Other requirements:** R v.4.1 or higher, Python v.2 or higher

**License:** GNU General Public License

**Any restrictions to use by non-academics:** A license is required to use the IBM solver which is only freely available for academics

## List of abbreviations

MoA: Mechanism of Action
TF: Transcription Factor
NES: Normalised Enrichment Score
NN: Nearest Neighbour

## Declarations

### Ethics approval and consent to participate

Not applicable

### Consent for publication

Not applicable

### Availability of data and materials

The dataset used in the case study is available from GEO (accession number GSE129254). The IBM Cplex optimiser is optional to use, and requires a license which can be obtained at https://www.ibm.com/products/ilog-cplex-optimization-studio, and is not included with the application or containers. The CBC solver is free to use without a license.

### Authors’ contributions

L.H.G developed the app in collaboration with R.H.B and wrote the manuscript, A.L. provided ideas for the app’s development and initiated the collaboration, H.B performed beta testing at Eli Lilly and Company, H.B, D.A.C and A.B supervised the project. All authors read and approved the final manuscript.

## Acknowledgements

The authors would like to thank Julio Saez-Rodriguez’s research group for beta testing the app and giving feedbac and Aurelien Dugourd for providing helper scripts for the application. We would also like to thank Jeff Kriske at Eli Lilly and Company for building the Singularity container solution.

